# The recent range expansion of Great Cormorants (*Phalacrocorax carbo hanedae*) in Hokkaido, Japan

**DOI:** 10.1101/2021.06.28.450098

**Authors:** Theodore E. Squires, Daisuke Aoki, Osamu Hasegawa

## Abstract

Though the Japanese subspecies of Great Cormorant (*Phalacrocorax carbo hanedae*) has increased its population in recent decades, information on its spread to the northern island of Hokkaido has not been reported outside of Japan. The purpose of this paper is to update the scientific community about the breeding and range ecology of *P. carbo*, and to provide comparative information on the abundant and similar looking resident Japanese Cormorant (*P. capillatus*). Several ornithological groups and researchers were contacted in order to gather information about the current distribution and breeding activities of *P. carbo* in the region. Here the findings of Japanese research groups, translated publications, and direct observations are made available.

## Introduction

During the initial phases of an environmental genetics study focusing on detecting Great Cormorants (*Phalacrocorax carbo hanedae*) in Hokkaido, Japan, it was discovered that no English information is published describing the status and distribution of *P. carbo* in the region. Despite this lack of available information, many Japanese researchers are aware of the spread of *P. carbo* on the north island and several breeding colonies have been identified. Here, recent evidence of the spread of *P. carbo* onto the island of Hokkaido is reviewed, and information from Japanese research groups, NPOs, and knowledgeable observers is consolidated.

*P. carbo* is generally broken into several subspecies with the local ‘hanedae’ historically occupying the length of the main Japanese island of Honshu south through Kyushu and the Ryuku Islands (Lethaby & Moores 1999, Clements 2007). Authors have also reported *P. carbo hanedae* as present in low numbers on the Korean peninsula where it is replaced by *P. carbo sinensis*, the dominate subspecies from southern China to north-central Europe wintering as far south as Indonesia (Lethaby and Moores, 1999; Clements, 2007). Other subspecies dominate globally through Africa, North-central Europe, Australia, New Zealand, and the east coast of North America to the Gulf of Mexico (BirdLife International^2^, 2018). On the main Japanese island of Honshu, *P. carbo* has been trained for centuries as a leashed diver for fishermen (Inoue-Maruyama *et al*. 2002). Across Japan, *P. carbo* populations have been expanding in both population and range significantly since the 1980s and are subjected to culling in some areas due to their predation on commercially important Sweetfish (*Plecoglossus altivelis*) among other species (Takahashi *et al*. 2006, Kumada *et al*. 2013, Natsumeda *et al*. 2014). This is in line with international trends for *P. carbo;* according to the IUCN, populations across Europe and North America appear to be increasing in both range and number with a total global estimate between 1.4 and 2.1 million individuals (BirdLife International^2^ 2018).

The cryptically similar Japanese Cormorant (*Phalacrocorax capillatus*; also referred to as Temmink’s Cormorant) is also seen throughout the Far East, though breeding is known to occur primarily on Hokkaido, Sakhalin, and the southern Kuril Islands, with smaller breeding groups seen on Honshu, the Korean Peninsula, and formerly Kyushu (Watanuki *et al*. 1988, Lethaby & Moores 1999). On the island of Hokkaido there are 77 recognized cities, townships, and villages along 2,676 km of coastline (Geographical Survey Institute, 2006). Several of these areas have multiple ports within their jurisdiction. The abundant *P. capillatus* regularly utilizes jetties surrounding these ports and offshore rocks for colony locations, so accurately mapping and estimating their population and distribution becomes difficult. In Japan, *P. capillatus* populations are estimated to be between 10,000 and 100,000 birds during the breeding season and global total population estimates are between 25,000 and 100,000, though no discrete estimate has been made for Hokkaido alone in recent years (BirdLife International^1^ 2018). Hokkaido census data identified approximately 1900 breeding pairs in the 1980s, and that number was believed to have increased in the preceding decade (Watanuki *et al*. 1988).

The most obvious differences in the two species’ breeding ecology is that *P. carbo* typically breeds near lakes and rivers where they forage, while *P. capillatus* tends to rely more heavily on costal and pelagic habitats where they have been observed to dive to significant depths for food (Lethaby & Moores 1999, Ishikawa & Watanuki 2002). *P. carbo* is also known to occasionally utilize port areas and offshore rocks for breeding in Japan. Though no research has been undertaken to evaluate interspecific interactions or competition between these species, we have provided general theories about their relationship in the discussion section.

While ornithologists and birders in Japan are generally aware of the presence of *P. carbo* on Hokkaido, many are not familiar with the details of their arrival. The range expansion has remained either unrecognized or unpublished by important multinational groups such as the IUCN and BirdLife International who do not show Hokkaido on current species range maps (IUCN 2014, BirdLife International^2^ 2018) and in recent guide books of both national and international scope (Clements 2007, Shimba 2007, Brazil 2009). This is surprising, as regular summer visits of *P. carbo* on the north island are known have started as early as the 1990s, and databases like eBird now show regular but debatable observations there year-round (eBird, 2018).

For this review, an attempt was made to communicate with as many researchers as possible, identify current colony locations, estimate a population size, and summarize Japanese publications to gain an understanding for the gradual increase of *P. carbo* in Hokkaido.

## Methods

### Study Area

Hokkaido is the 2^nd^ largest and northernmost island of Japan. It is approximately the size of Austria at 83,454 km^2^, taking up about 22% of the landmass of the Japanese archipelago. Despite its large size, the human population density is disproportionately low compared to the rest of Japan, and the island is comprised mostly of forested and agricultural lands. Rivers run throughout the island fed by abundant central mountains and access to some mountainous areas can be difficult due to heavy snowfalls from October through April.

### Inquiries

Intensive formal inquiries started in mid-October of 2015 and ended in early February of 2016. Several authors were contacted to create a network of individuals capable of providing firsthand knowledge of the birds or access to relevant Japanese publications. Eventually some companies and research groups were contacted to access information unavailable to the English-speaking community. Because the status of *P. capillatus* is well established on Hokkaido, all inquiries asked if the correspondents knew the locations of *P. carbo* colonies specifically, and often extended to include questions of population numbers, foraging habits, and site fidelity. Most inquiries were sent in both Japanese and English and assistance in translating responses was sought when necessary. Japanese research papers regarding *P. carbo* on Hokkaido were gradually translated as time permitted. It was assumed that all who were contacted would be capable of distinguishing *P. carbo* from *P. capillatus*.

### Independent Observations

From March 2016 to March 2017, trips were taken to visit several colony and roosting sites after their locations were elucidated. Point counts were taken at water sampling points on over 110 occasions with the aid of spotting scopes, digital cameras, and binoculars. Birds were positively identified as *P. carbo* based on the guidelines provided by Lethaby and Moores’ 1999 publication. Flight and foraging behaviors are nearly identical between *P. carbo* and *P. capillatus*, so we particularly relied on the shape of the gape line for identification, which protrudes backwards giving a sharper appearance in *P. capillatus* while in *P. carbo* the rear of this line appears more flush with rounded skin behind the lower mandible. The tail is shorter and overall size is smaller in *P. capillatus* than in *P. carbo*, but at a great distance these field marks can be impossible to distinguish and even up close it may take a trained eye to correctly determine species. In addition to field marks, habitat and location were used as an initial consideration.

## Results

Thirty-six of the contacted individuals responded to our inquiries leading to information for approximately 1800 *P. carbo* nests on Hokkaido. Near the town of Horonobe around 1500 nests were reported, though that colony changed location and was reduced significantly from the preceding year. Our personal observations estimated less than 600 nests in the new Horonobe location. Approximately 400 nests were observed east of Kimoma Wetland near the town of Sarufutsu. An additional 100 nests were reported to be present in the port of Abashiri. A colony of 50 nests near the town of Tsukigata was observed to disperse in April of 2016, while other groups of similar size were reported in the ports of Esashi, Motoinabu, and Monbestu. The inland areas of Wakubetsu and Setana were also believed to host smaller colonies of 25 individuals or less. In addition to these colonial nesting sites, the presence of stable roosting areas was confirmed south of Asahikawa upstream from Kamui Kotan, and near the south bank of the mouth of the Tokachi River. One other nesting or roosting area was reported to possibly exist near the town of Nanae. These findings are clearly summarized in Table 1 (also see Fig. 1). Based on collected information, almost all *P. carbo* leave Hokkaido during the cold winter months, and no birds have been recorded on the island during February. Migrating *P. carbo* generally arrive in March and leave in November. The overall population estimate for breeding *P. carbo* on Hokkaido in 2016 was at least 2,500 individual birds but probably not exceeding 6,500 individuals. The total number of non-breeding *P. carbo* and newly recruited birds has been theorized to be between 2,000 to as many as 5,500 individuals, leading to population estimates as high as 12,000 individuals on the island by the end of the 2016 breeding season.

**Figure 1.**
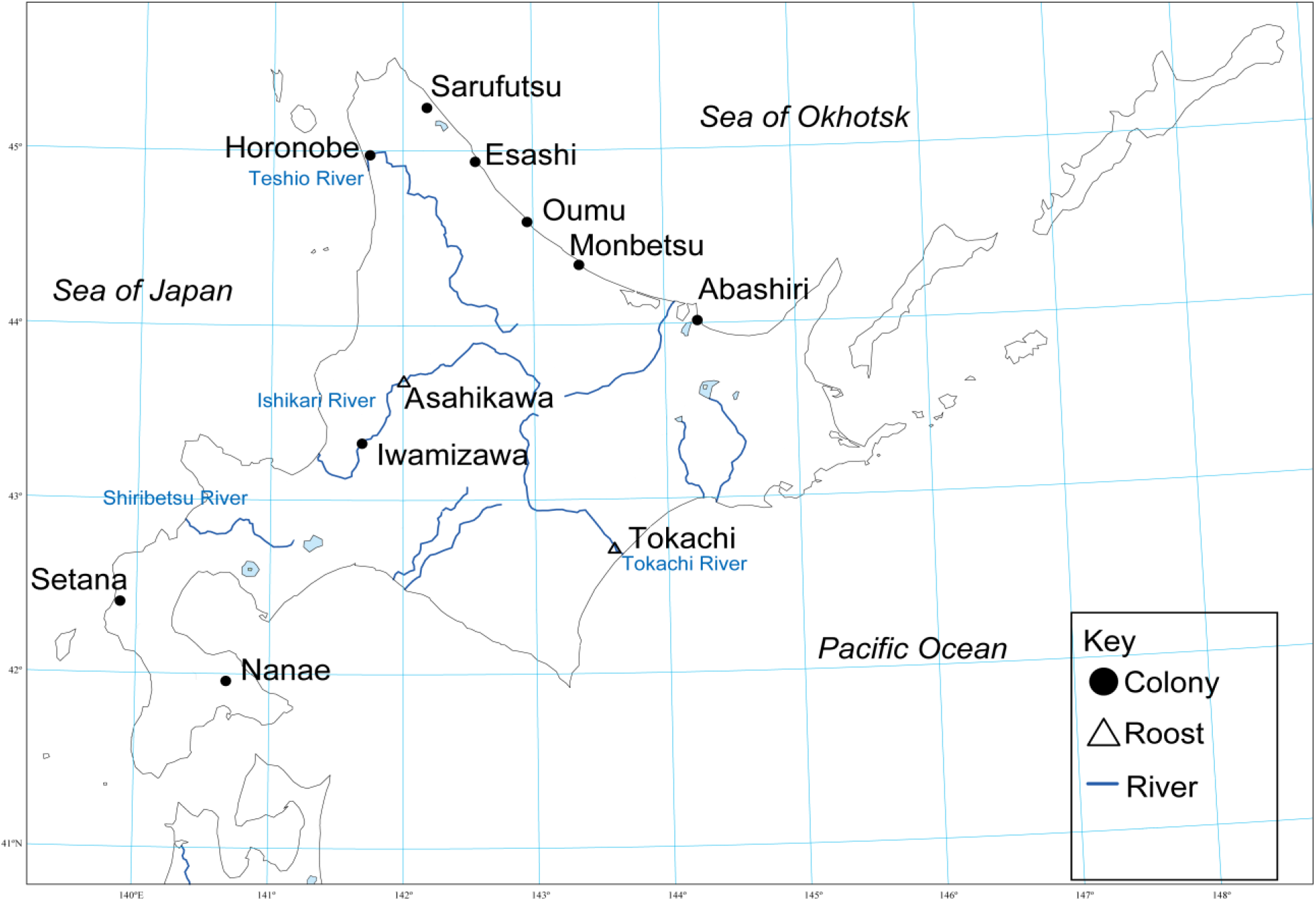
Infographic showing the distribution of *P. carbo* colonies on Hokkaido during our studies in 2016.

**Table 1.** Information about Hokkaido’s *P. carbo* nesting colonies and roosting distribution based on Kazuhiro Odate’s unpublished summary of HGCRG findings for 2016 and his personal correspondence, and additional roosting information provided by Kentaro Takagi. Original findings by the HGCRG and provided by Takagi are highlighted in grey while direct observations and additions from the authors are shown without shading.

| Area of Hokkaido | Detail of area | Lat. N ; Lon. E (Decimal Degrees) | Nests | Note |
| --- | --- | --- | --- | --- |
| Horonobe | Confluence of Sarobetsu and Teshio River | 44.97; 141.74 | 600 | Moved in 2015 for unknown reasons, previously 1500 nests located west side of Penke Marsh |
| Sarufutsu | Kimoma wetland | 45.28; 142.19 | 400 |  |
| Esashi | Esashi port | 44.55; 142.35 | 50 |  |
| Oumu | Motoinabu port | 44.36; 142.57 | 50 | Disappeared 2016 |
| Monbetsu | Monbetsu port | 44.20; 143.21 | 50 |  |
| Wakubetsu | Ichigousen Bridge left side | 44.13; 143.36 | 25 | Founded 2016 |
| Abashiri | Abashiri port | 44.01; 144.17 | 100 |  |
| Iwamizawa | East-southeast of Tsukigata Town on the Ishikari R. east bank | 43.33; 141.69 | 50 | Dispersed April 2016 |
| Setana | Makomanai dam | 42.49; 139.91 | ? | Unconfirmed |
| Nanae | Junsai Wetland | 41.99; 140.63 | ? | Disappeared prior to 2015? Roost only? |
| Asahikawa | Vicinity of Kamui Kotan | 43.73; 142.20 | 0 | Roost |
| Tokachi | Northwest of Otsusaiwai | 42.71; 143.61 | 0 | Roost |

## Discussion

Based on translated research, visits of *P. carbo* to Hokkaido were first recorded in April, June, and October of 1974 near the city of Nemuro (Fujimaki, 2007). Small numbers of *P. carbo* were subsequently observed in 1979, 1982, 1990, 1991, 1994, and 1995 though it was at the time considered a rare incidental species (Fujimaki, 2007). The migratory behavior of breeding colonies was recognized in the northernmost part of Honshu on the Shimokita Peninsula in Aomori prefecture by the 1980s (Abe, 2005; Kato, 2008). *P. carbo* then became a regular visitor after flocks of up to 100 birds were observed at the confluence of the Shinotsu and Ishikari Rivers on Hokkaido in April of 1999 (Kato, 2008). In 2001, the very first breeding reports of at least 16 *P. carbo* nests were verified on Hokkaido in the vicinities of Horonobe and Nanae (Hokkaido Wild Birds PR Dept., 2002; Fujimaki, 2007; Kato, 2008). In general the population is understood to have been increasing since that time. Representative Yasuhiro Kawasaki from the Wild Bird Society of Japan (WBSJ) in Okhotsk echoed many informal conversations and indicated there had been a general increase in *P. carbo* sightings on Hokkaido since the year 2000.

We believe *P. carbo* does not overwinter on Hokkaido due to the complete freezing of almost all inland lakes that they rely on, and range maps showing migratory mainland Asian populations generally wintering below 40° north (Shimba, 2007; BirdLife International^2^, 2018). Similarly, older Japanese publications noted a few individual birds present in the winter months, but suggested that there is a significant outmigration and concluded that *P. carbo* would be unlikely to survive the midwinter average −8°C degrees, responsible for freezing their preferred habitats (Fujimaki, 2007). On Hokkaido, most sightings appear to taper off by mid-November and resume in early March and any stragglers would likely move towards the southern coasts of the island; further we found no verified instances of *P. carbo* presence on the island in February. Within Japan, several genetically distinct subpopulations of *P. carbo* have been identified, and it appears that Hokkaido’s birds actually are more closely related to *P. carbo* populations sampled in central western Japan (Hasegawa *et al*., 2007; Great Cormorant Protection and Management Report, 2012). Despite the genetic history, there is no evidence to suggest wintering areas for *P. carbo* migrating from Hokkaido and this would make an interesting future study.

Our 2016 total population estimate for *P. carbo* on Hokkaido was placed between 4,500 and 12,000 individual birds, of which we directly observed about 1,300 individuals. Port colonies were observed to nest on jetties and offshore rocks while inland colonies reliably nested in trees within 100 meters of rivers or large ponds. No clear preference for foliage type was observed inland as both conifers and deciduous trees were used for nesting. Instances of colony failure or abandonment could not be definitively explained, but theories included harassment by large birds of prey, racoon dog (*Nyctereutes procyonoides viverrinus*), and incidental disturbances from nearby farming activities. Non-breeding (foraging) groups of up to 70 individuals regularly perched on gravel bars, tree limbs, exposed rocks, and in one case a large group rested on a construction device above the Tsukigata dam.

On Hokkaido, *P. capillatus* is generally recognized to be dominant in all coastal areas while *P. carbo* better utilizes inland rivers and lakes. No research has been undertaken to see if this follows any spatial patterns related to bathymetry, but it appears that colonies of *P. carbo* have arisen more at Hokkaido’s northern ports along the shallow Okhotsk Sea than in other frequently deeper seaside areas. Our personal observations suggest that *P. carbo* numbers across Hokkaido’s coasts are still dwarfed by abundant *P. capillatus* colonies from which many individuals remain foraging year-round. This discrepancy aids in field identification though both species can be seen on coastlines and far inland while foraging (Lethaby and Moores, 1999; See. Fig 2). There is no evidence to suggest that *P. carbo* numbers have negatively influenced P. capillatus where they share spaces, and instead it appears that both species populations have increased in Hokkaido in recent decades. Because *P. carbo* is a frequent riverine forager throughout its range, it may have superior performance in shallower waters than *P. capillatus*. Further, *P. capillatus* being primarily an ocean bird, likely performs better when foraging at depth in tidal conditions. In the future, foraging studies in the areas where these species frequently cohabit may determine if they are competing for similar food resources or experiencing more success. Understanding the many factors that could be influencing P. carbo’s range expansion with will require significant expansion of research into the species, its interspecific interactions, and the regional environment.

**Figure 2.**
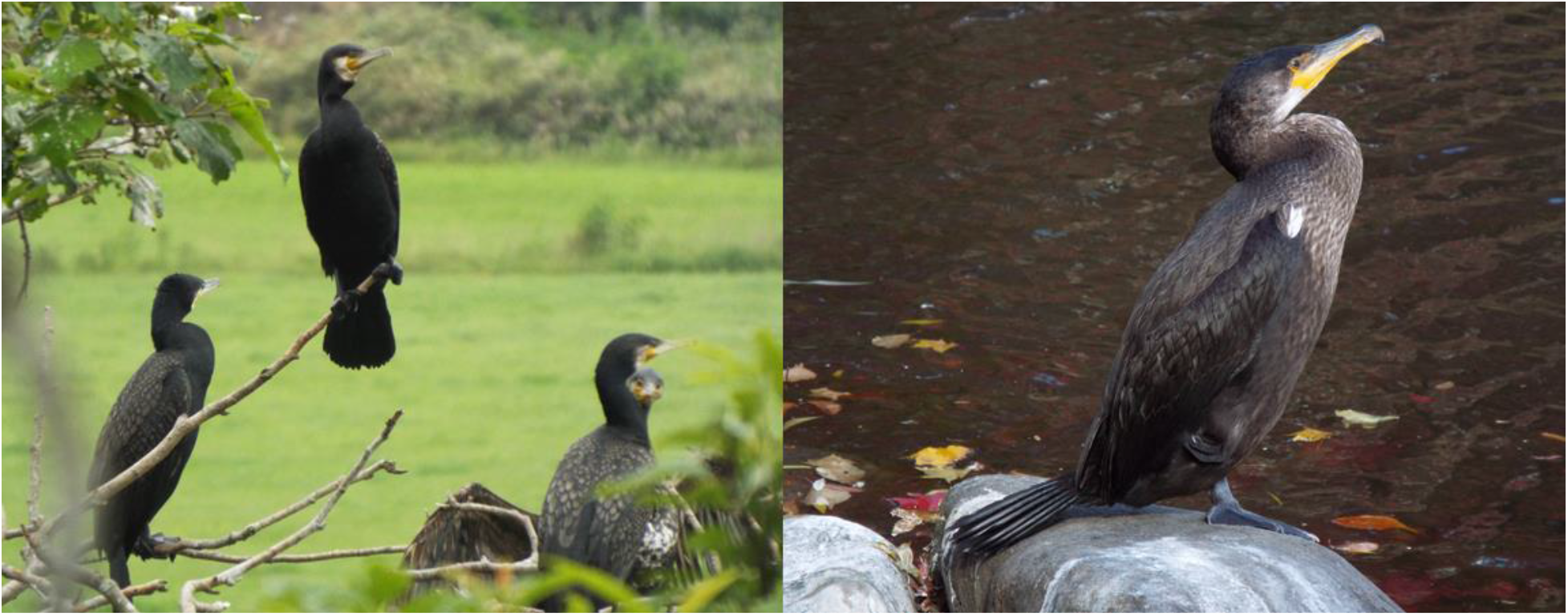
Side by side pictures of *P. Carbo* (left) and *P. Capillatus* (right) for comparison. Photograph of *P. carbo* taken in May of 2016 at the Sarufutsu colony site. Photograph of *P. capillatus* taken in early November 2016 shows an individual 17 km inland at Nakajima Park in Sapporo City.

Our intensive search for information left the impression that there are few immediate concerns about the commercial impacts of increasing populations of *P. carbo* on Hokkaido. Our formal findings are in direct contrast to reports from individual e-bird users who have reported *P. carbo* present on Hokkaido through the winter months (eBird, 2017). The presence of *P. capillatus* in the area and their similar appearance to *P. carbo* likely explains this situation. Because the expansion of *P. carbo* is not a significant topic of investigation domestically, it is not surprising that details of *P. carbo’s* life history in Japan has only been sparsely addressed in English publications. Hopefully, future research on the *hanedae* subspecies can be brought to the attention of a wider audience and the information provided here aids in holistically understanding the migration and breeding of *P. carbo* on Hokkaido. For this work, many people who were contacted could only speculate or generalize details about arrival and departure times, population estimates, and dietary information, so we believe that further inquiries will continue to form an integral part of ongoing research.

## Acknowledgements

Many thanks to the members of Hokkaido Great Cormorant Research Group (HGCRG), especially Kazuhiro Odate, for allowing us to share their findings. Sincere thanks to two anonymous reviewers who greatly improved the quality of our publication with their suggestions. Regards to Kentaro Takagi, Yasuko Suzuki, Kayoko Kameda, and all Hokkaido chapters of WBSJ for sharing information that helped to build a bigger picture. Additional recognition is deserved for Yuzo Fujimaki for putting us in touch with most of our contacts. Lastly, thanks to members of the Animal Ecology lab at Hokkaido University’s Department of Agriculture, who provided skilled and professional translations upon request.

## References

BirdLife International^1^. 2018. Species factsheet: *Phalacrocorax capillatus*. Available online at: http://datazone.birdlife.org/species/factsheet/22696799

BirdLife International^2^. 2018. Species factsheet: *Phalacrocorax carbo*. Available online at: http://datazone.birdlife.org/species/factsheet/22696792

Brazil M.A. 2009. Birds of East Asia: China, Taiwan, Korea, Japan, and Russia. Princeton University Press, Princeton.

Clements J. F. 2007. The Clements Checklist of Birds of the World, 6th Edition. Cornell University Press, Ithaca.

eBird. 2018. eBird: An online database of bird distribution and abundance [web application]. eBird, Cornell Lab of Ornithology, Ithaca, New York. Available online at: http://www.ebird.org

Fujimaki Y. 2007. Recent Distribution of Cormorant *Phalacrocorax carbo* and Rook *Corvus frugilegus* in Hokkaido, northern Japan. Strix 25: 109–119. [In Japanese]

Geographical Survey Institute. 2006. Islands, area and length of coastline of national land. Ministry of Land, Infrastructure and Transport. Japan Coast Guard. Available online at: http://tochi.mlit.go.jp/english/data-concerning-lands/outline-of-national-land

Great Cormorant Protection and Management Report. 2012. Kawau no hogo kanri ni kansuru repōto [Report about the protection and management of Great Cormorants (2012 version)]. Japanese Ministry of the Environment. [In Japanese]

Hasegawa, O., Ishigaki M., Fukuda M., Nīdzuma Y. & Azuma S. 2007. Kyūgekina bunpu kakudai no katei de, kawau no iden-teki kōzō wa dō keisei sa reta ka? (Happyō yōshi) [Has the Great Cormorant formed genetic structures following rapid spread? (announcement summary)]. Ornithological Society of Japan, Abstracts for the Annual Meeting No. 123. [In Japanese]

Hokkaido Wild Birds Public Relations Department. 2002. Shinbun jōhō kara - kawau eisō dōnai hatsu kakunin [Information from newspaper- first confirmation of Great Cormorants nesting in Hokkaido]. NPO Bird Research: Bird News from Hokkaido No. 129. [In Japanese]

Inoue-Maruyama M., Ueda Y., Yamashita T., Nishida-Umehara C., Matsuda Y., Masegi T. & Ito S. 2002. Molecular sexing of Japanese cormorants used for traditional fishing on the Nagara River in Gifu City. Animal Sci. Journal 73: 417–420.

Ishikawa K. & Watanuki Y. 2002. Sex and individual differences in foraging behavior of Japanese cormorants in years of different prey availability. Journal of Ethology 20:49–54.

IUCN Redlist. 2014. BirdLife International and NatureServe (2014) Bird Species Distribution Maps of the World. *Phalacrocorax carbo*. The IUCN Red List of Threatened Species. Version 2016-1. Available online at: http://maps.iucnredlist.org/map.html?id=22696792

Kumada N., Tomoko A., Tsuboi J., Akihiko A. & Fujioka M. 2013. The multi-scale aggregative response of cormorants to the mass stocking of fish in rivers. Fish Sci. 137: 81–87.

Lethaby N. & Moores N. 1999. Identification of Temminck’s Cormorant. Dutch Birding 21:1–8

Kato N. 2008. Hokkaidō ni okeru kawau no bunpu kakudai [Distribution expansion of the Great Cormorant in Hokkaido] NPO Bird Research: Bird News from Hokkaido No. 154. [In Japanese]

Natsumeda T., Sakano H., Tsuruta T., Kameda K. & Iguchi K. 2015. Immigration of the common cormorant Phalacrocorax carbo hanedae into inland areas of the northern part of Nagano Prefecture, eastern Japan, inferred from stable isotopes of carbon, nitrogen and Sulphur in rivers. Fish Sci. 81:131–137.

Shimba T. 2007. Photographic Guide to Birds of Japan and Northeast Asia. Yale University Press, New Haven.

Takahashi T., Kameda K., Kawamura M. & Nakajima T. 2006. Food habits of great cormorant *Phalacrocorax carbo hanedae* at Lake Biwa, Japan, with special reference to ayu *Plecoglossus altivelis altivelis*. Fishe Sci. 72: 477–484.

Abe S. 2005. Heisei 16-nendo Heisei 17-nendo kawau seisoku chōsa [Fiscal Year 2004 - 2005 Great Cormorant Habitat Survey]. Mutsu City Cultural Property Survey Report No. 34: 117–124. [In Japanese]

Watanuki Y., Kondo N. & Nakagawa H. 1988. Status of Seabirds breeding in Hokkaido. Japanese Journal of Ornithology 37: 17–32. [In Japanese]

